# Four assessment methods to measure student gains in a graduate course on mathematical modeling in cell biology

**DOI:** 10.1101/346890

**Authors:** Renee Dale, Naohiro Kato, Bill Wischusen

## Abstract

The push for mathematics and quantitative skill development in biology is greater than ever before. The perceptions of these techniques are a barrier to their implementation on a grand scale. Methods to assess the efficacy of approaches to teaching quantitative skills are needed to determine if they effectively overcome social stereotypes and fears of math. In our project we study the effect of a mathematical modeling course in cell biology. We developed four assessments to understand if hands-on modeling increases student understanding of the concepts while simultaneously reducing fear of math. Two concept inventories track students’ learning gains both in general and specific knowledge to the course content. Two surveys quantify students’ quantitative confidence and opinions of the usefulness of these techniques in biology. We discuss the effectiveness of our methods and the implications for assessment development in mathematical biology education.

## 1. Introduction

Mathematics and computational skills are increasingly critical in the field of biology. Whether a student is looking to work in public health, education, or basic research, new computational techniques will be required to deal with the increasingly large datasets obtained thanks to electronic and high-throughput data-gathering methods.

*Vision and Change* calls for more quantitative training in the biological sciences [4]. The social perception of math being for ‘geniuses’ serves to enhance the struggles of disadvantaged groups [1, 2, 3]. Educators experiencing math anxiety are not confident in their ability to teach students the quantitative skills necessary for a technological age [6]. When biology students are exposed to new methods involving math they often ‘freak out’ [6]. This is partially due to social perceptions of math and to belief that new methods are inherently more difficult [3, 1, 2]. Research has shown that as students engage in studying math and technology their attitudes improve [5, 9, 7]. These experiences have also led to increased efficacy in their studies and engagement when learning math in context of their existing interests [8, 10]. However, post-secondary education is often not up to the challenge [6]. Students actively transfer out of college degree programs to avoid mathematics. Math anxiety or gender bias present in teachers may prevent them from demonstrating the utility or importance of mathematics to their students prior to choosing their majors in college.

Given these issues in quantitative biology skills, the ability to understand and quantify the effectiveness of an exercise or course is increasingly important. We developed 4 methods to measure the changes in our students. To measure student learning gains over the semester, we developed a general biology concept inventory and two concept inventories specific to course content. We developed a survey to measure student attitudinal changes toward mathematics and quantitative biology. Since validation of the effectiveness of a survey to accurately measure as intended, we further developed an open-ended survey. We find that our assessment methods perform well and discuss their implications.

## 2. Methods

### 2.1. Study Population

Our study population was a group of 7 graduate students studying engineering or biology. The students were taking a course in mathematical modeling in cell biology. We received IRB approval for this study (#E10547).

### 2.2. Opinion and confidence survey

We developed a 5-point Likert scale survey to capture student’s attitudes on topics related to mathematical modeling. The survey had four categories: techniques, experimental design, knowledge, and confidence. We masked the true intentions of our survey by including four question categories with other, related questions included. This is to avoid ‘social desirability’ response bias, since students are aware that mathematical biologists would be viewing their responses. Survey respondents tend to respond to seem more socially desirable [20]. This tendency is of direct concern to us, due to the social perceptions of math and programming. Our primary interest was the confidence of students in performing quantitative techniques such as programming and math, and their opinion to the importance or utility of these methods as wet-lab researchers in their fields. These questions were scattered about the 4 categories.

### 2.3. Concept inventory

We curated a set of 24 concept inventory questions on general biological knowledge from the literature [11, 12, 13, 14, 15, 16, 17, 18, 19]. As the course consisted of 4 general biological topics (enzymes, gene expression, diffusion, and signaling), we selected a set of 6 questions per course topic.

### 2.4. Specific inventory

Since many of the concept inventory questions we found contained information from introductory biology courses, which was many years ago for our study population of graduate students, we developed two specific concept inventory surveys based on specific modeling case studies: the g-protein signaling pathway and firefly luciferase enzymatic reactions. Students took the pre-test immediately before applying their modeling skills on the case study, and post-test after.

### 2.5. Open-ended opinion survey

We designed the open-ended opinion survey to validate our previous surveys. The open-ended survey included questions about math confidence, utility, and student opinion of the value of the course. All questions are available in the supplemental materials.

### 2.5. Analysis

To quantify gains in learning, confidence, or opinion we normalized for values greater than zero using the equation 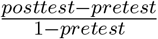 [21]. We used a cutoff of zero since we assume that a negative learning gain represents student guessing. Negative learning gains were observed fairly commonly in the concept inventories. For the opinion and confidence survey, only one question by one respondent dropped by one point in the post-survey. We considered this to be a non-significant zero change.

## 3. Results

The course discussed general programming techniques and mathematical concepts, followed by modeling of 4 general biological concepts - enzymes, gene expression, diffusion, and signaling. This was followed by two modeling case studies, the g-protein signaling pathway and firefly luciferase enzymatic reactions. To determine how the course affected student perceptions related to mathematical modeling in biology, we developed the 4 assessment methods illustrated in Fig. 1.

**Figure 1.**
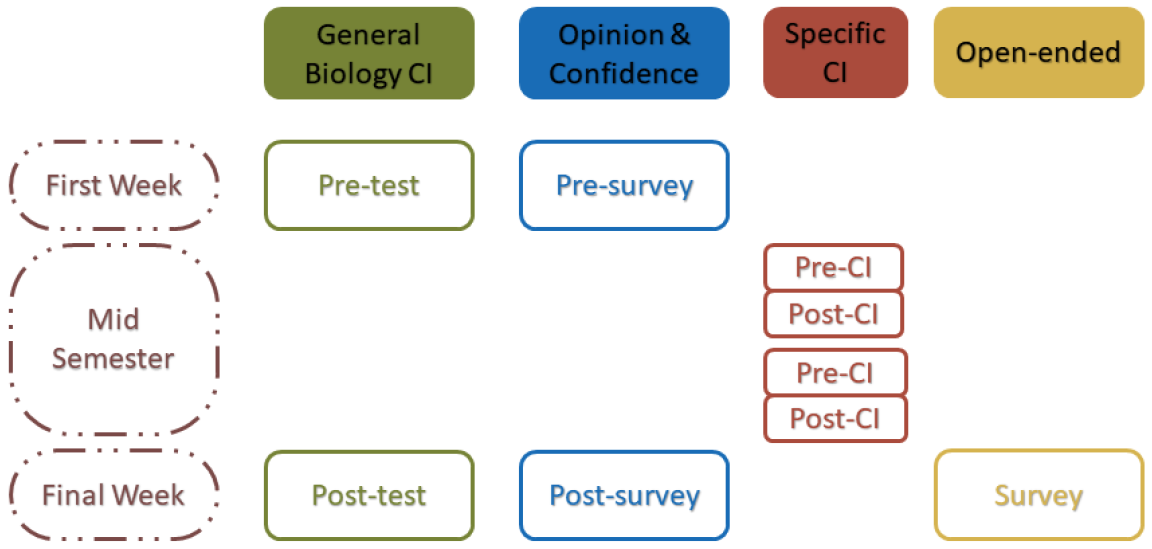
Student gains were measured by the four assessment methods spread out over the semester.

**Figure 2.**
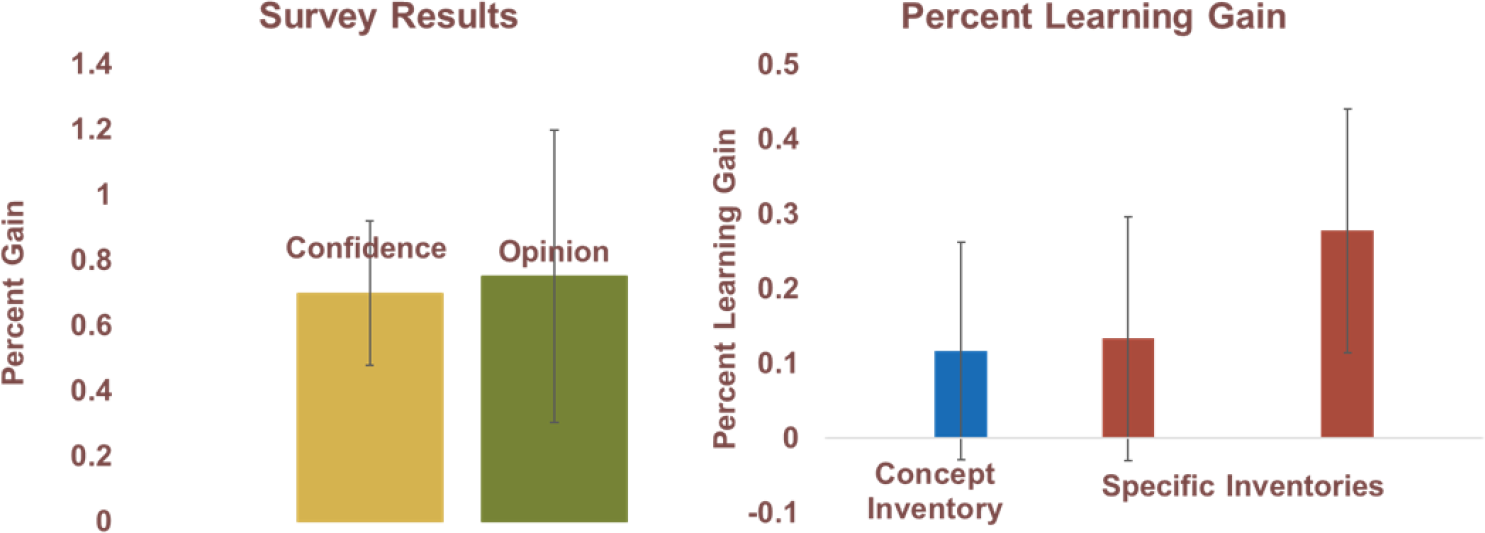
Outcomes of a mathematical modeling course measured using three methods. The students reported large gains in their math and programming confidence (*mean = 0.70*) and their opinion of the importance and utility of quantitative methods in biology (*mean = 0.75*). Our methods also measured learning gains in a general biology concept inventory (*mean = 0.15*) as well as learning gains specific to two modeling scenarios (*mean = 0.13, 0.28*).

Student learning gains as measured by the general biology concept inventory averaged 15% from the pre-test to the post-test. Our two concept inventories covering material explicitly covered during the course showed larger learning gains. The two specific concept inventories covered the g-protein signaling pathway, an average gain of 13%, and the firefly luciferase enzyme kinetics showed an average gain of 28%.

The survey intended to measure student confidence in quantitative skill and opinion of the usefulness of those skills showed much larger gains. Our survey was constructed with 4 apparent categories. The survey questions that were related to our research question were distributed throughout the 4 categories. This was intended to avoid response bias, as students might feel pressured to appear more confident [20]. Our survey showed a 70% improvement in student confidence, and 75% improvement in opinion of the utility of quantitative techniques in biology.

The open-ended survey included questions that were redundant with the survey intended to understand confidence and opinion changes. Other respondents answered positively, and generally in agreement with the observed effects. In answer to the question “Do you see yourself modeling in the future?” a student answered: *I can see myself modeling in the future to determine what can be the best initial conditions in a biological experiments.*

We found the most interesting and promising answers to the question “Do you think this course will have helped you form interdisciplinary collaboration in the future?” *Yes, it has showed how important and helpful a mathematical model can be to a biological system.* The final question “What were your previous methods to understand or explain a biological system in an educational or publication context? Do you think this will change after you have done some mathematical modeling?”

> *Wet lab, trial and error. I will use mathematics for a better planification of experiments*
>
> *I would dig in literature and take online courses on introductory levels of systems biology.*
>
> *Previous methods only included understanding steps in a diagram. Now, one must consider quantities and speed and each step.*

## 4. Discussion

Our results indicate that having graduate students apply novel quantitative methods to topics related to their fields improves their confidence in quantitative methods as well as their opinion of the usefulness of those methods. This result was captured in both our opinion survey and the open-ended survey intended to validate it. Our concept inventories did not show strong learning gains, but we believe this is due to question formulation. Survey respondents misunderstanding a question is another possible issue in assessment development [**?**]. In the open-ended survey, we noticed at least one incidence of misunderstood question. In answer to the question, “Do you think experimental design is affected by knowledge of mathematical modeling?”, a student answered: *No. I don’t think modeling can overpower the benefits of experiments on any day. Each is needed in its own way. Once you learn about the connection, it will be affected and possibly improved.* The student interpreted this question as an forcing a choice between either experimental or mathematical methods, whereas our intention was to understand how the students might apply their new skills in realistic research scenarios.

Another possible issue is that the general biology concept inventory was too general, and the specific inventories were too specific given the time constraints for a single modeling application task. Overall, we feel this indicates the importance of multiple measures to understand the effectiveness of tasks involving new quantitative skills. Although the learning gains of the students as measured by our methods in our study were not large, the students reported strong affinity to apply the tasks that they learned. This may be due to a distinction between fully understanding or being engaged in the biological system, and the technical skill required to carry out a quantitative analysis of that system.

